# Rate of functional network maturity and the role of environmental factors

**DOI:** 10.1101/2025.09.05.674401

**Authors:** B Hughes-Small, S Saljoughi, RA Stevenson, A Soddu, B Stojanoski

## Abstract

The developmental period from childhood to adolescence is marked by significant changes to brain function that support higher-level cognition. Environmental factors such as socioeconomic status and adversity can have an outsized but distinct influences on neurocognitive development. Here, we examined the differential influences of SES (i.e., parental education and neighbourhood safety) and adversity on the maturation rate of functional networks in children and adolescents. Using resting-state fMRI data, independent component analysis with dual regression was computed to identify six networks (Default Mode Network (DMN), Left Executive Control Network (ECN), Right ECN, Hippocampal (HPC), Salience, and Sensorimotor networks) of interest in children and adolescents aged 7 to 15 (*N*=216, acquired from the Healthy Brain Network). A neural maturity index was generated based on the degree of similarity between the spatial configuration of each networks in every youth brain to that of a sex-matched adult template. Regression analyses were used to determine the association between neural maturity, environmental factors, and social-cognitive abilities. We found one sensory (sensorimotor) and two association (default mode and executive control) networks matured faster than other networks. Only the rate of maturity of the DMN and HPC were associated with environmental factors. Maturity of the DMN was associated with less adversity and better social cognitive ability, whereas maturity of the HPC network was associated with younger participants with higher IQs. Moreover, these effects were stronger in females than males. Our results highlight the importance of examining the unique contributions of distinct dimensions of childhood environments on neurocognitive development.

## Introduction

The transition from childhood to adolescence signifies a period of tremendous social and cognitive development. In addition to blossoming cognitive abilities such as reasoning and working memory, the brain undergoes massive structural and functional changes during this period (Fox et al., 2010). Crucially, the environment that children grow up in can have an outsized influence on brain development and the cognitive processes they support (Tooley et al., 2021). For instance, positive environments that are safe, stimulating and contain strong social relationships support healthy brain development (Hackman & Farah, 2009; Leonard et al., 2019). Conversely, a negative environment that contains stressors, such as low socioeconomic status (i.e., limited access to resources), concerns over neighbourhood safety (Vargas, Rakesh, & McLaughlin, 2025), and exposure to adversity can hinder neurocognitive development (Brieant et al., 2021; Chahal et al., 2022; Duncan et al., 2012; Evans, 2016; Kim-Spoon et al., 2021; Rakesh et al., 2023a; Rakesh et al., 2025).

Early life adversity and socioeconomic status (SES) are related in important ways and reflect environmental characteristics children have no control over. However, they are not inexorably linked; children from low-SES environments are not guaranteed to encounter adverse experiences, and children reared in high-SES backgrounds are not excluded from the possibility of experiencing adversity. However, SES and childhood adversity are also not completely independent. For example, a shared characteristic of low-SES environments and those high in adversity is the limited access to resources; unavailable resources can lead to the presence of adverse experiences such as stress and unmet developmental needs (Walsh et al., 2019). Despite the complex interplay and partial overlap between SES and adversity, their unique contributions to cognitive and brain development remain unclear.

Low-SES and adversity can have deleterious effects on cognitive development. For instance, children raised in low SES environments perform worse on measures of reasoning, inhibition, and working memory (Burneo-Garces et al., 2018; Finn et al., 2017; Ming et al., 2021) and, in general, have poorer executive functioning compared to children raised in high SES environments (Korous et al., 2020; Lawson et al., 2018; Rakesh et al., 2025; Yu et al., 2024). Similarly, children who have experienced forms of maltreatment perform more poorly on various cognitive tasks. For instance, victims of sexual and physical abuse, neglect and those exposed to unsafe and disadvantaged neighbourhoods demonstrate poorer memory for verbal and numerical information (Irigaray et al., 2013; McCrory et al., 2010; Shonkoff & Garner, 2012; Taylor et al., 2020; Yu et al., 2024) and often have global deficits in executive function (Dodaj et al., 2017; Nikulina & Widom, 2013; Majer et al., 2010).

Childhood socioeconomic status and adversity not only influence the development of cognition but also the development of the brain, in particular, the rate of brain maturation (Rakesh et al., 2023b; Roubinov Meaney, & Boyce, 2021). Like its influence on cognition, the influence of SES on brain development is multi-dimensional and not uniform (Tooley et al., 2021), resulting in seemingly contradictory findings. For instance, children raised in low SES backgrounds have been shown to have both increased (more mature) and decreased (less mature) connectivity between primary and higher-order brain networks (Rakesh et al., 2025; Rakesh et al., 2021a,b; Ramphal et al., 2020; Tooley et al., 2020; 2021), whereas children from higher-income backgrounds tend to have more mature network configuration (Gao et al., 2015a; Tooley et al., 2020, 2021).

Early life adversity also does not appear to have a uniform impact on network maturation (Lindenmuth et al., 2025). In general, more adverse experiences are often associated with accelerated maturation (Tooley et al., 2021). For example, young children exposed to high-adversity backgrounds compared to children in low-adversity spaces demonstrate accelerated maturation in the form of structure-function coupling (i.e., correlation between structural and functional connectivity; Chan et al., 2024). Likewise, childhood adversity is linked to increased connectivity within various regions of the DMN (Holz et al., 2023), a hallmark of network segregation and brain maturation (Fair et al., 2007, 2009; Grayson & Fair, 2017; Satterthwaite et al., 2013). However, there is evidence that adversity in the form of psychosocial stress is associated with patterns of reduced connectivity within the frontoparietal network (Holz et al., 2023), a characteristic most prominent in younger brains. Likewise, increased (maturity) and decreased (developing) connectivity within the salience network have been associated with childhood adversity (McLaughlin et al., 2019).

Given these mixed findings, the exact nature in which SES influences functional network development remains poorly understood. The findings suggest that SES and ELA do not impact brain development along a single dimension, and different cortical networks may have unique sensitivities and reactions to environmental influences, ultimately contributing to differences in cognition (Rakesh et al., 2021a,b; Ramphal et al., 2020; Tooley et al., 2021; Yu et al., 2024).

The aim of the current study was to improve our understanding of the influence of socioeconomic status and early life adversity on functional network maturation and social and cognitive abilities. Using resting-state and cognitive measures from the Healthy Brain Network, we used independent component analysis with dual regression along with a template matching procedure with adult brains to explore whether 1) functional brain networks mature at different rates, 2) socioeconomic status and early life adversity have distinct influences on different networks across the brain, 3) there is a relationship between cognitive abilities, environmental influences, and network maturity, and 4) environmental influences brain networks are sex-specific. We hypothesized that sensory networks, such as the sensorimotor network, would be more adult-like than association networks, such as the DMN and FPN. Additionally, we hypothesized that early life adversity, more than socioeconomic status, would be associated with network maturation, and that this association would be related to cognitive functioning in a sex-dependent manner.

## Materials and methods

### Participants and Data Acquisition

Data was retrieved and analyzed from the Health Brain Network (HBN) Biobank, a Child Mind Institute initiative. The HBN is an ongoing initiative to collect behavioural, physiological, clinical (diagnostic consultation), genetic and neuroimaging data from individuals aged 5 to 21 (described in Alexander et al., 2017), to understand mental health and wellness. This study was approved by the Chesapeake Institutional Review Board and the Research Ethics Board at Ontario Tech University. More information regarding the dataset can be found at http://fcon_1000.projects.nitrc.org/indi/cmi_healthy_brain_network/. In total, 216 (127 male) participants aged 7 to 15 were acquired from HBN releases 1 through 10. Participants were included in the current study if 1) both anatomical and resting state functional MRI data were available, 2) the data passed quality assurance threshold (see below), 3) all environmental and 4) cognitive measures were completed (with an IQ greater than 66 limiting cognitive impairments that might interfere with study procedure; Alexander et al., 2017).

### Adult template

The adult template was generated using multi-site data exclusively from healthy participants acquired across several established repositories, such as the Human Connectome Project (HCP 900), ABIDE, and the AOMIC collection. The template was generated using functional MRI (1,243 subjects; mean age 27), and T1-weighted imaging (1,665 subjects; mean age 33). Separate templates were generated based on sex, allowing for sex-specific comparisons with the youth sample.

### Environmental and Cognitive Variables

Parental education was assessed through the Barratt Simplified Measure of Social Status, which captures the mean parental education on a Likert scale, ranging from 7^th^ grade to graduate education (see Table 1). The PhenX Neighbourhood Safety Scale, which measures parent-rated neighbourhood safety, were also measured on a Likert Scale. Responses ranged from “1 = Strongly Agree” to “5 = Strongly Disagree,” with higher values indicating less neighbourhood safety. The Negative Life Events Scale provided a measure of parent-reported total number of adverse experiences based on a 21-item scale (Sandler et al. 1991). The scale includes questions about stressful or upsetting experiences in a child’s life, particularly those involving family conflict, illness or injury, emotional distress, and major changes or losses. It covers events such as parental illness, arguments, job loss, death of loved ones, and disruptions like changing schools or losing a friend. Cognitive abilities were measured using the Wechsler Intelligence Scale for Children-V (WISC), a measure of higher-order cognitive abilities, and dimensions of social cognition were assessed using the Social Responsivity Scale-2 (SRS), an Autism Spectrum Disorder (ASD) screening tool (Constantino et al., 2003; Wechsler, 2014). Finally, age, sex, and race were included as demographic variables.

**Table 1.** Summary of participant demographics.

| Variable | Category | Mean (SD)/N(%) | Range |
| --- | --- | --- | --- |
| Age, years |  | 10.9 (2.5) | 7.01–15.9 |
| Negative Life Events |  | 7.1 (3.4) | 0–17 |
| Neighbourhood Safety |  | 2.2 (1.2) | 1–4 |
| Cognitive Measures |  |  |  |
|  | Fluid Reasoning (WISC) | 104.8 (15.5) | 67–140 |
|  | Processing Speed (WISC) | 97.4 (16.0) | 49–155 |
|  | Verbal Comprehension (WISC) | 106.8 (15.2) | 65–150 |
|  | Visual Spatial Ability (WISC) | 102.4 (16.7) | 57–144 |
|  | Working Memory (WISC) | 103.2 (15.6) | 62–146 |
|  | Full-Scale IQ (WISC) | 103.8 (16.1) | 63–141 |
|  | Social Cognition (SRS) | 7.7 (5.6) | 0–27 |
| Parental Education |  | 18.3 (2.8) | 3–21 |
|  | Less than 7th grade | 1 (0.5%) | 3 – 5.9 |
|  | Junior high/Middle school, 9th grade | 1 (0.5%) | 6–8.9 |
|  | Partial high school, 10th or 11th grade | 4 (1.8%) | 9–11.9 |
|  | High school graduate | 9 (4.2%) | 12–14.9 |
|  | Partial college, at least one year | 38 (17.6%) | 15–17.9 |
|  | College/University education | 102 (47.2%) | 18–20.9 |
|  | Graduate degree | 61 (28.2%) | 21 |
| Race |  |  |  |
|  | White/Caucasian | 101 (46.8%) |  |
|  | Black/African American | 42 (19.4%) |  |
|  | Hispanic | 16 (7.4%) |  |
|  | Asian | 6 (2.8%) |  |
|  | Indian | 2 (0.9%) |  |
|  | Multiracial (2+) | 36 (16.7%) |  |
|  | Other | 4 (1.8%) |  |
|  | Did not specify/answer | 9 (4.2%) |  |

### Preprocessing and analysis rs-fMRI Preprocessing

Neuroimaging data included T1-weighted anatomical MRI scans and functional MRI data acquisition from 5-minute resting-state sessions. Patients were instructed to open and close their eyes at various times during each fMRI session. A gradient-echo planar imaging pulse sequence (TR = 800 ms, TE = 30 ms, Flip Angle = 31 degrees, whole brain coverage 60 slices, resolution 2.4 × 2.4 mm2) was used to collect functional imaging, and structural imaging was acquired using a high-resolution T1-weighted MPRAGE in 224 sagittal (TR = 2500 ms, TE = 3.15 ms, resolution 0.8 × 0.8 mm2).

Resting-state fMRI data were preprocessed and analyzed in native space using GraphICA, a Brainet-developed (https://www.brainet.ca) platform based on FSL 6.0.3 (Jenkinson et al., 2012). T1-weighted anatomic images included the following preprocessing steps: bias field correction, brain extraction, tissue-type segmentation (cerebrospinal fluid, gray matter, white matter) and subcortical segmentation.

Conversely, functional data preprocessing included skull stripping, section-timing correction, spatial smoothing (ceiling of 1.5 voxel size), high-pass filtering of 100 seconds, independent component analysis (ICA)-based Automatic Removal Of Motion Artifacts (AROMA) and Nuisance regressors which included motion correction based on six motion parameters (x, y, z, roll, pitch, yaw), white matter and cerebrospinal fluid.

GraphICA uses displacement quality control when processing the fMRI scans using the following equation:

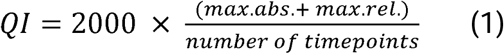

Where *max.abs.* represents the maximum absolute displacement of a volume within a reference time point, and *max.rel.* is the maximum relative displacement of a volume within the subsequent time point. Quality index (QI) captures abrupt and gradual changes in head movement or position (Ribeiro de Paula et al., 2017) and is used to identify poor-quality data. Participant data that reported a quality index (QI) measure exceeding 50 were excluded from the sample.

### Functional Networks

GraphICA performs Independent Component Analysis (ICA) with dual regression implemented in FSL (Jenkinson et al., 2012). The dual regression unfolds in two stages. First, the inputs to the first stage include preprocessed fMRI data for individual participants, along with the group-level spatial (independent component) template maps, which are used to perform spatial regression to extract subject-specific time series for each independent component map. In the second stage, the subject-specific time courses from the first stage, along with the subject’s fMRI data, are used to conduct a temporal regression to extract subject-specific spatial maps for each independent component map. The output of this process is a set of independent component maps generated for each network; dual regression was implemented to identify subject-specific spatial maps using 11 resting-state network masks, covering functional connectivity across the entire brain. The networks included the default mode network (DMN), executive control network left (ECNL), executive control network right (ECNR), hippocampal network (HPC), language network, salience network, auditory network, sensorimotor network (SenMot), visual lateral network, visual medial network and visual occipital network.

For the purposes of the current study, we focused on six independent component maps: DMN, ECNL, ECNR, HPC, salience network, and SenMot network (Figure 1; adapted from Halder, 2023). The ICA maps formed the basis for comparing our youth sample to a normative database comprising 1243 healthy adults (433 females, 28 ± 17 and 810 male adults, 27 ± 16), who act as a control group to which all nodes are compared.

**Figure 1.**
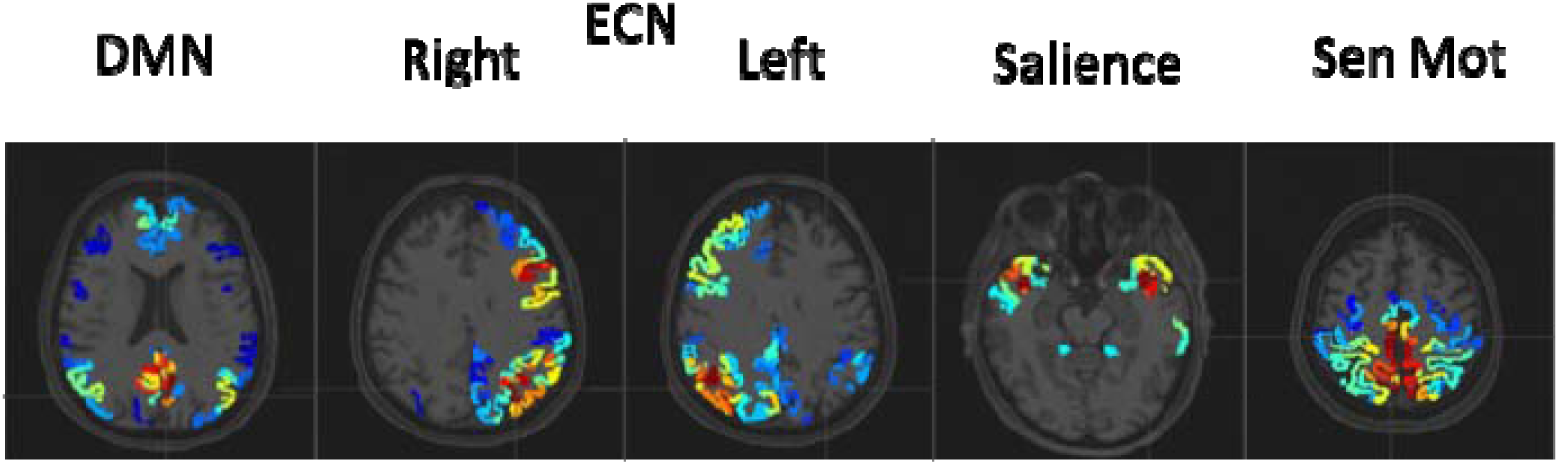
Note. Representation of cortial location of each network. Networks were extracted using group-ICA and superimposed on a structural T1 image. From left to right: default mode network, right and left executive control network, salience network, sensorimotor network, overlaid on T1 scan corresponding to adult template.

### Regional Parcellation

Structural data (each participant’s T1-weighted image) were automatically segmented within the GraphICA pipeline using FreeSurfer (v7.1.0). Functional data were further parcellated through GraphICA’s automated gradient-weighted Markov Random Field model procedure (Schaefer et al., 2018). The parcellation model described (Schaefer et al., 2018) incorporates three key components that balance different segmentation goals: a global similarity term that groups together brain areas with similar image intensities, a local gradient term that detects abrupt changes in functional connectivity between neighbouring brain locations, and a spatial connectedness term that ensures regions remain spatially coherent. This approach resulted in 832 distinct brain parcels, which were subsequently used as network nodes to generate functional networks of interest (see below). The parcellation framework (Schaefer et al., 2018) provides a method for dividing the brain (cerebral cortex) into regions using intrinsic functional connectivity that aims to achieve a balance between detailed local information and broad global coverage.

### Network Construction

After the functional resting-state networks were co-registered at the individual participant level (using ICA with dual regression), mean z-score statistics at the network level were generated by taking the average z-score values for all voxels comprising the 832 parcels (regions of interest). That is, z-standardized correlation coefficients quantify the contribution of each nodal time series to its corresponding resting-state network. To mitigate the influence of noise and exclude spurious or low-magnitude z-values, thresholding of the z-scores (statistical maps) was performed using a data-driven approach for network matrices based on mixture modelling (Churchill et al., 2016) at the individual subject level. This method leverages fractional moment series of blood oxygen level-dependent (BOLD) signal distributions to enhance the robustness of effective connectivity analyses (Bielczyk et al., 2019). By minimizing noise and removing unreliable connections, the procedure improves the interpretability of functional network structure, emphasizing biologically meaningful interactions and increasing cross-dataset replicability, particularly when addressing the challenges posed by non-Gaussian BOLD signal characteristics.

### Measures of Neural Maturity

Using the thresholded Z-score maps, we computed a neural maturity index based on a distance correlation value reflecting the degree of similarity between the individual and the adult template (1243 healthy adults). The distance correlation measures the strength of the relationship between two variables. The advantage over Pearson’s correlation is that distance correlation can detect non-linear relationships and work with multi-dimensional data. Distance correlation (dCor) was computed for each participant (*X*) and template network (η) using Equation 2. A value of 0 represents no correlation, and a value of 1 represents maximum correlation; the higher the correlation value, the more similar the subject is to the template database, and therefore, the child or adolescent brain is considered more adult-like.

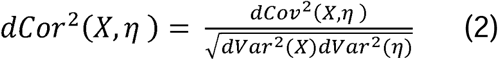

### Statistical Analysis

All statistical analyses were conducted using R Studio. Neural maturity was subjected to a one-way repeated ANOVA to identify significant differences in network maturation across the networks of interest. Linear regression analyses were employed to determine which environmental factors and cognitive abilities were most strongly associated with the functional maturation in the six networks of interest. Parental education (SES), NLES (Adversity), neighbourhood safety, age, sex, social cognition (SRS) and higher-order cognitive measures (WISC) were included to predict neural maturity across networks of interest. To account for potential confounding effects of residual head motion, mean framewise displacement (FD) was included as a covariate of no interest in all regression models (Power et al. 2012). That a linear regression with all predictors of interest were conducted for each network of interest. False Discovery Rate (FDR) corrections were applied to correct for multiple comparisons (q = 0.05). To replicate previous findings, pearson correlations (df = 214) were conducted between all cognitive (fluid reasoning, processing speed, social cognition, verbal comprehension, visual spatial ability and working memory) and environmental measures (parental education, adversity, and neighbourhood safety), and independent samples t-tests (df = 214) was conducted to compare the cognitive measures between the two sex groups (see Supplementary Materials for results).

## Results

### Demographic and Environmental Factors

The final sample included 127 (59%) males and 89 (41%) females. Table 1 contains the average scores and standard deviations for our demographic variables (age, parental education, negative life events and neighbourhood safety); five cognitive abilities (i.e., fluid reasoning, processing speed, verbal comprehension, visual spatial ability and working memory) and Full-Scale IQ based on the Wechsler Intelligence Scale for Children (WISC); as well as Social Cognitive ability based on the Social Responsivity Scale (SRS).

### Sex and Cognitive Ability

All cognitive performance measures were compared between the sexes. We failed to find any difference between males and females for any of the cognitive measures (t _(208.5)_ < −1.81; p > 0.25), except for processing speed. We found that females (M_PSI_ = 100.8) had faster processing speeds than males (M_PSI_ = 95; t _(208.45)_ = 2.75, p = 0.042).

### Neural Maturity Across Networks

We used a neural maturity index (NMI) across the six networks of interest to identify which networks matured more rapidly than others. We found that the bilateral executive control networks (ECN) matured most rapidly relative to the other regions (M_ECNL_ = 0.65, SD_ECNL_ = 0.65; M_ECNR_ 0.66, SD_ECNR_ = 0.09). The sensorimotor network (M = 0.61, SD = 0.15) and default mode network (DMN) (M = 0.56, SD = 0.13) matured more rapidly than the salience (M = 0.42, SD = 0.12) and hippocampal networks (M = 0.28, SD = 0.13), which reported the slowest rate of maturity (the least similarity to the adult template).

One-way repeated ANOVA revealed a main effect if network; that is, at least one network was significantly different than another (F_(5, 1290)_ = 312.55, p < 0.001). Post hoc analyses revealed the following results (df = 430). The DMN was significantly less mature than the ECNL (t = −8.27, p < 0.001, Cohen’s d = −0.80), ECNR (t = −9.46, p < 0.001, Cohen’s d = −0.91) and SenMot (t = −4.37, p <0.001, Cohen’s d = −0.42), and more mature than the HPC (t = 21.31, p < 0.001, Cohen’s d = 2.05) and salience (t = 11.27, p < 0.001, Cohen’s d = 1.08). The ECNL was more mature than the HPC (t = 31.28, p < 0.001, Cohen’s d = 3.01), salience (t = 21.01, p < 0.001, Cohen’s d = 2.02) and SenMot (t = 2.82, p = 0.005, Cohen’s d = 0.27), but was not significantly different from the ECNR (t = −0.95; p = 0.34, Cohen’s d = −0.092). The ECNR was more mature than the HPC (t = 33.34, p < 0.001, Cohen’s d = 3.21), salience (t = 22.84, p < 0.001, Cohen’s d = 2.2) and SenMot (t = 3.68, p = 0.003, Cohen’s d = 0.35). The HPC was less mature than the salience (t = −10.95, p < 0.001, Cohen’s d = −1.05) and SenMot (t = − 24.05, p < 0.001, Cohen’s d = −2.31). Finally, the salience was less mature than the SenMot (t = −14.86, p < 0.001, Cohen’s d = −1.43). See Figure 2 for visual representations.

**Figure 2.**
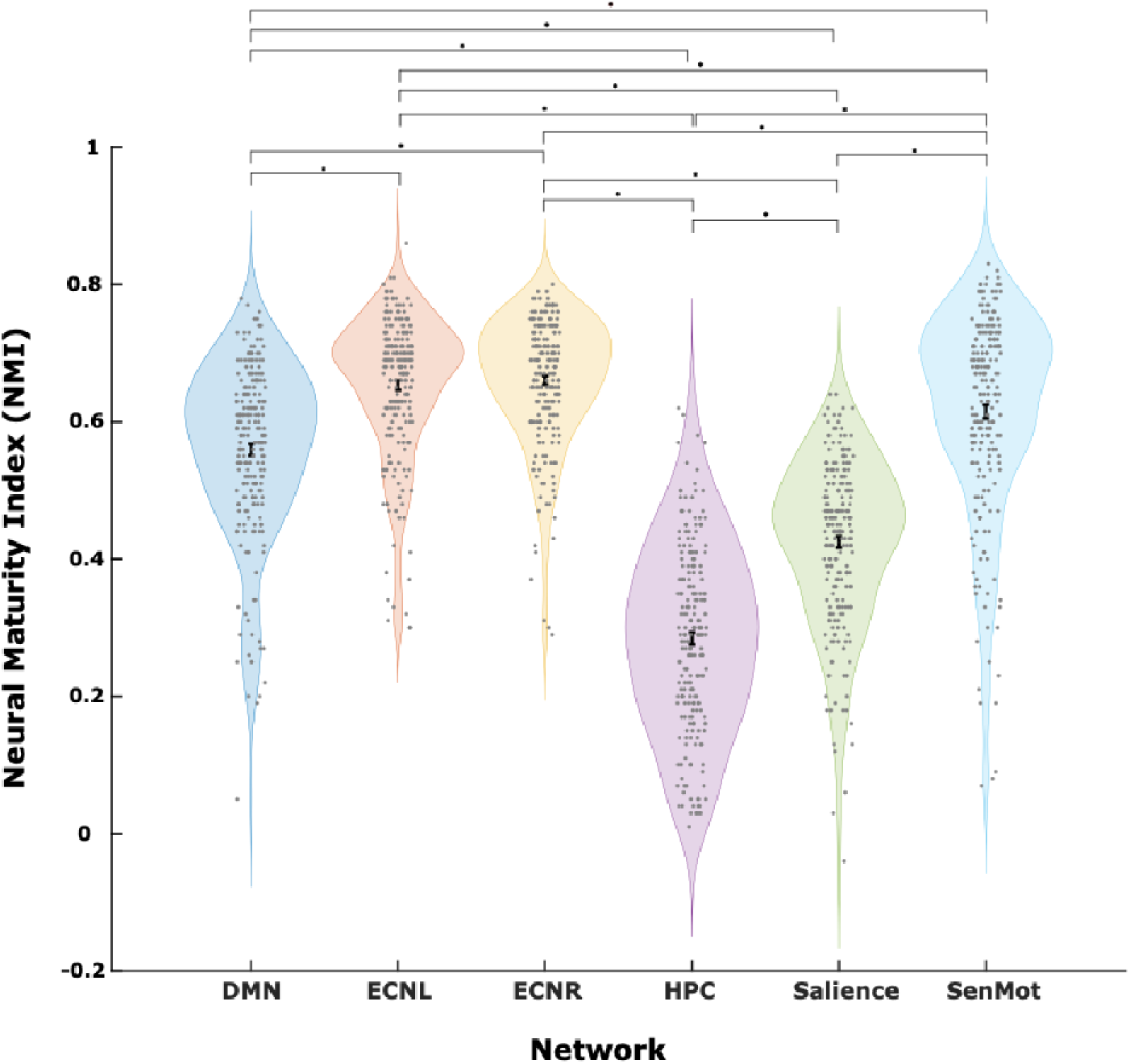
Neural Maturity Across the Six Networks of Interest Note. Violin plots representing the distribution of Neural Maturity Index values across the six regions of interest (colour-coded) with standard error of the mean (black line) and a scatter of each data point (gray). Lines and asterisks indicate networks where the difference in NMI was statistically significant. DMN = Default Mode Network. ECNL = Executive Control Network Left. ECNR = Executive Control Network Right. HPC = Hippocampal. SenMot = Sensory Motor Network.

### Regression Analyses Default Mode Network

Of the six networks explored, only two reported significant results and survived multiple corrections, one of which was the default mode network. Specifically, we found sex, social cognition, and adverse events (total) significantly predicted the functional maturity of the DMN (F _(8, 204)_ = 7.69, p < 0.001). More precisely, female participants (t = 4.88, p < 0.001), those with better social cognitive abilities (t = −2.29, p = 0.02), and those with fewer adverse events (t = −2.36, p = 0.018) reported more adult-looking functional connectivity of the DMN (see Figure 3). These variables accounted for 23.18% (Adjusted R^2^ = 20.17 %) of the variance.

**Figure 3.**
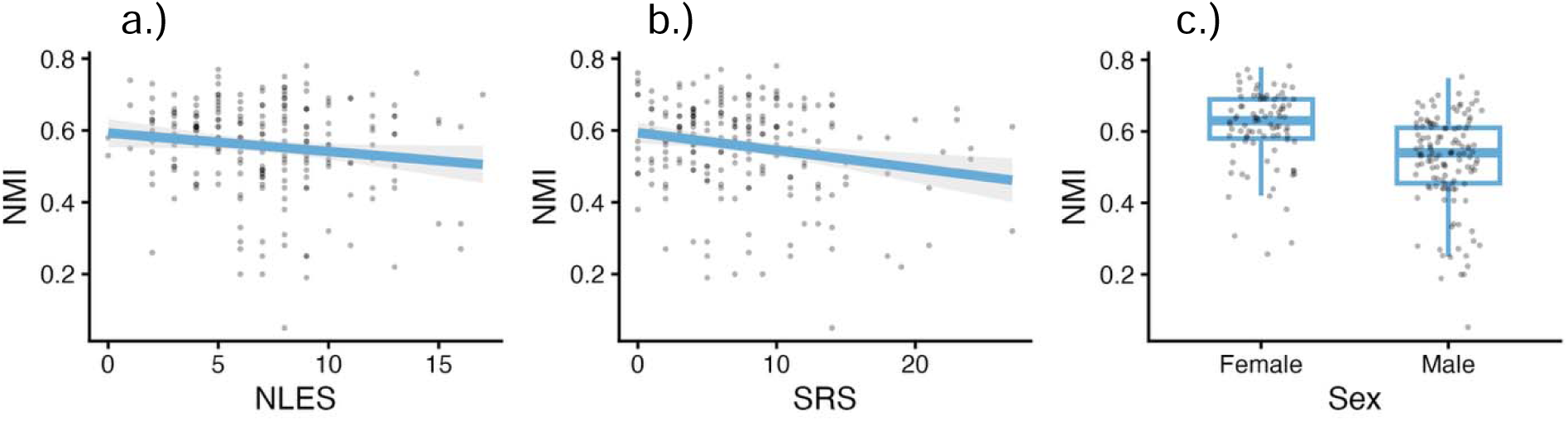
Relationship between neural maturity of the default mode network and social cognition and negative life events. *Note.* Visual distributions of the variables that significantly predicted neural maturity within the default mode network (DMN). a.) Scatter plot of DMN maturity across the number of adverse experiences. b.) Scatter plot of DMN maturity across social cognition Scores. For figures *a* and *b*, points represent participants, and blue lines represent the regression line of best fit. c.) Box plot of DMN maturity between the sexes. Points represent participants.

### Hippocampal Network

The second network to show significant results was the hippocampal network. Within this network, full-scale intelligence quotient (FSIQ), age and sex significantly predicted neural maturity (F _(8, 204)_ = 4.59, p < 0.001). More specifically, those with higher IQ (t = 1.94, p = 0.05), younger participants (t = −3.84, p < 0.001), and females (t = 3.78, p < 0.001) reported more similarity to the adult template for this network (see Figure 4), accounting for 15.25 % (Adjusted R^2^ = 11.93 %) of the variance within the model.

**Figure 4.**
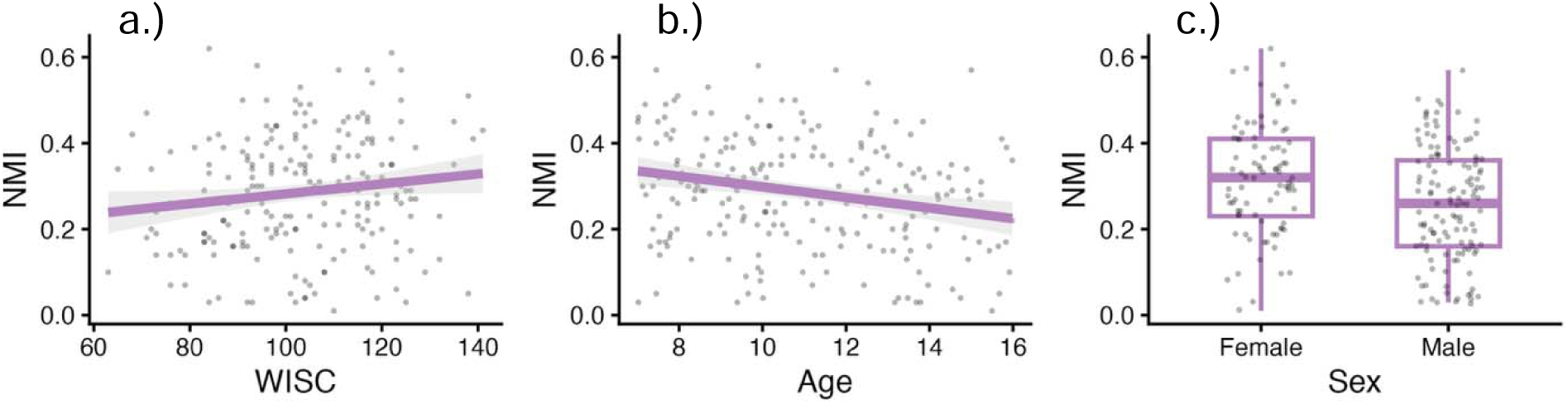
Predicting neural maturity in the hippocampal network Note. Visual distributions of the variables that significantly predicted neural maturity within the hippocampal network (HPC). a.) Scatter plot of HPC maturity across Age. b.) Scatter plot of HPC maturity across FSIQ scores. For figures *a* and *b*, points represent participants, and blue lines represent the regression line of best fit. c.) Box plot of HPC maturity between the sexes. Points represent participants.

## Discussion

We investigated the rate of brain maturation across several functional networks of interest, and the influence of environmental factors, such as socioeconomic status and early life adversity, on functional network maturity, executive function and social cognition in children and adolescents. We found significant differences in the degree of network maturity; the sensorimotor network, along with two association networks (the default mode and the bilateral executive control networks), showed similar degrees of maturity and were the most adult-like. Importantly, differences in the rate of network maturation were sex-specific, with more mature networks in females relative to males. Moreover, we found that parental education and adverse experiences were differentially associated with neurocognitive development; neural maturity was not related to parental education or neighbourhood safety, but parental education and neighbourhood safety were significantly related to different aspects of cognition, whereas neural maturity was associated with adversity (a less adult-like default mode network was linked to more negative experiences) and social cognitive ability.

Executive function and social cognitive ability were significantly associated with parental education. Specifically, children and adolescents with more educated parents reported better performance on all executive function measures (fluid reasoning, processing speed, verbal comprehension, visual-spatial ability and working memory), full-scale intelligence quotient (IQ) (a composite measure of the listed abilities), and social cognitive ability. Parental education, one dimension of socioeconomic status, is exceptionally influential on cognitive development across the lifespan (Halse et al., 2019; Kaplan et al., 2001; Rindermann & Ceci, 2018; Roberts et al., 1999; Sha et al., 2018). One mechanism explaining the link between parental education and better cognitive performance is that more educated parents provide more enriching environments for their children, which, in turn, supports greater cognitive development (Rakesh et al., 2024). Indeed, greater socioeconomic status has been associated with differences in parenting styles and techniques (Davis-Kean et al., 2019, 2021; Freeberg & Payne, 1967) and more enriching home environments (Davis-Kean, 2005; Rosen et al., 2020; Votruba-Drzal, 2003), which positively contribute to cognitive development.

Neighbourhoods rated as safer were also associated with better performance on social cognition and all higher-order cognitive measures. Neighbourhood-level characteristics that may be contributing to cognitive development include resource availability (Cloney et al., 2015; Moore et al., 2008) and the presence of green spaces (Flouri et al., 2019; Reuben et al., 2019), both of which are important for cognitive functioning (Li, Whittle, & Rakesh, 2025). Conversely, this result may also reflect an indirect relationship with parental education, as parents with more education rated their neighbourhoods as safer. The finding that neighbourhood safety is associated with cognition further illustrates the influential power the environment has on cognition. Still, future research is needed to clarify the nature of the relationship between neighbourhood safety, socioeconomic status and cognition.

Although there is evidence for the association between adversity and poorer cognitive performance, especially on memory measures (Irigaray et al., 2013; Shonkoff & Garner, 2012; Yu et al., 2024), none of our cognitive measures, including working memory, correlated with adversity. One potential explanation for this finding is that our sample included many highly-educated parents (nearly half had a college or university degree), and parental education offers a protective influence that moderates the impact of adversity. Indeed, higher parental education (among other SES factors) has been associated with positive parenting practices and enriched environments, which are also associated with better cognitive performance (Davis-Kean et al., 2019, 2021; Freeberg & Payne, 1967; Rakesh et al., 2024; Rosen et al., 2020). Therefore, not only do highly-educated parents create an environment that fosters cognitive development, but they may also buffer the negative effects of adversity on cognitive functioning (Beck et al., 2025; Crouch et al., 2018; McLaughlin et al., 2014; Roubinov & Boyce, 2017).

We found females demonstrated significantly higher scores on the processing speed subscale of the WISC relative to males. In fact, this was the only sex-based difference across the different socio-cognitive measures. This result replicates previous research that across the lifespan females outperform males in various measures of processing speed (Camarata & Woodcock, 2006; Roivainen, 2011). However, there is also evidence for a male advantage on visual tasks and a female advantage on verbal tasks (Herlitz et al., 2013; Hyde & Linn, 1988; Voyer et al., 1995), and we were surprised not to find similar sex-based differences. This pattern of results may be accounted for by the bias towards highly-educated parents in our sample which are obscuring potential sex differences, and this pattern of results may not be consistent across SES groups.

We found the greatest maturity in the bilateral executive control networks (ECN), followed by the sensorimotor (SenMot), default mode network (DMN), salience and the hippocampal (HPC) network was the least adult like. These results indicate that resting-state cortical networks follow distinct developmental trajectories that result in different degrees of maturity during childhood and adolescence.

That we found adult-like topographies in the only sensory network of interest aligns with the sensory-association axis of neurocognitive development that argues sensory networks develop sooner than higher-order networks, such as the ECN and DMN (Sydnor et al., 2021, 2023). Indeed, adult-like topography of the sensorimotor network has been found in infants (Edde et al., 2021), and this network exhibits fewer changes after birth than other networks (Gao et al., 2015a, 2015b). Interestingly, we also found adult-like architecture in left and right ECNs and the DMN (at levels comparable to the SenMot network), a result that seems incompatible with classic models that argue higher-order networks develop later than sensory networks (Edde et al., 2021; Gao et al., 2015a, 2015b; Sydnor et al., 2021). However, this result aligns with recent work demonstrating that both ends of the sensory-association axis appear to develop simultaneously rather than progressing linearly from sensory to association regions (Dong et al., 2021; Tooley et al., 2022). This new framework of network maturation is consistent with evidence for adult-like forms of the DMN in utero, as well as strong age-dependence on parcel-level functional connectivity within frontal regions of the DMN and the visual networks (Tooley et al., 2022). It also aligns with gradient measures of network segregation in prefrontal regions, resembling sensory networks by adolescence (Baum et al., 2020), and with metrics of network maturity for the control network and DMN, which peak around 11.5 and 12.5 years, respectively (Dong et al., 2021). However, not all association networks mature at the same rate; we found the salience network was less adult like than the ECN and DMN and falls at an intermediate stage along the neural maturation continuum (Dong et al., 2021). This suggests the salience network may follow a more protracted developmental trajectory, potentially reflecting its role in integrating information across multiple domains and supporting the emerging capacity for flexible cognitive control during adolescence (Luna, 2009; Krönke et al., 2020; Menon & Uddin, 2010).

We found the rate of maturity in the DMN was associated with better social cognitive abilities and fewer adverse experiences, and this relationship was stronger in female than male participants. Importantly, this pattern of results was independent of age.

Given there were no age effects, the relationship between adversity and neural maturity of the DMN does not appear to depend on an accumulation of adverse experiences in adolescents relative to children. Instead, it suggests a potential dose effect on this network’s maturity rate; more adversity experiences were associated with a less mature DMN. This result runs counter to our prediction that adversity would be associated with accelerated rates of network maturity, which has been previously reported (Callaghan & Tottenham, 2016). One potential reason for our result is that not all adverse experiences have the same effect on the brain. The NLES contains items that are related to trauma and aggression, which are linked to older-looking brains, but also contains items related to emotional distress which is often associated with younger-looking brains (Beck et al., 2025). Moreover, the DMN was the only network to demonstrate a relationship with adversity, raising important questions about the unique sensitivities of this network to the environment and how that may impact the development of cognitive abilities associated with the DMN.

We also found that better social cognitive ability was associated with greater DMN maturity. A once-believed dormant network has since been repeatedly connected to self-reflective thought, theory of mind and other social cognition (Mars et al., 2012; Schilbach et al., 2008; Xie et al., 2016). A more mature DMN likely supports more sophisticated social cognitive abilities, a finding that is consistent with the idea that better social cognition is associated with greater within-DMN connectivity, a hallmark of cortical development (Fox et al., 2017; Che et al., 2014; Washington & VanMeter, 2015), whereas reduced connectivity of the DMN has been associated with deficits in social cognition (Dodell-Feder et al., 2014). It is important to note that the causal direction of the relationship is unclear; it is possible that greater maturity of the DMN supports better social cognition, but it is also possible that better social cognition facilitates greater network maturation. Future studies are needed to identify the nature and direction of the relationship between social cognition skills and DMN maturity.

Although we failed to find environmental factors significantly associated with hippocampal network maturity, we did find a relationship between neural maturity, age, sex and cognition in this network. That is, greater IQ, females and younger participants had greater hippocampal network maturity.

Specifically, individuals with higher IQ also had a more mature HPC network. This result suggests that the HPC is associated with higher-order functioning and that more mature HPC networks appear to be related to better overall cognitive functioning. Indeed, greater network efficiency is associated with higher intelligence (Li et al., 2009; van den Heuvel et al., 2009), and increased hippocampal and prefrontal cortex (both ventral and medial regions) connectivity predicted problem-solving, future planning (Calabro et al., 2020; Li et al., 2020), and reasoning (Li et al., 2020), all higher-order cognitive abilities (Diamond, 2013) that comprise the full-scale IQ (Wechsler, 2014). One potential explanation for our finding is that more adult-like HPC networks are functionally more efficient (Supekar et al., 2009), and this efficiency gives rise to better cognition and higher IQ. However, future research is needed to better understand the relationship between network efficiency and cognitive development.

One of the more intriguing results of this study was that younger children demonstrated greater similarity to the adult HPC template than did adolescents. A result that may indicate that hippocampal network maturity may not follow a linear trajectory.

Although the literature on the development of the functional hippocampal network is sparse (Qin et al., 2014), insights from the structural literature may help interpret this pattern of results. For example, previous research examining the structure of the hippocampal network has found a curvilinear developmental progression for the head of the hippocampus (Canada et al., 2020; Tamnes et al., 2018). Including a larger age range in future research could uncover the nature of potential non-linear developmental trends in the HPC from childhood to adolescence.

Despite previous literature that has found associations between parental education and brain development (Noble et al., 2015; Cermakova et al., 2023), to our surprise, we did not find a connection between parental education and neural maturity across any of the networks of interest. One potential reason we failed to find this relationship is that the specific measure of education we used may not directly impact the rate at which networks mature. Relatedly, parental education is only one aspect of socioeconomic status, broadly understood as the various factors in our lives that create or diminish access to resources (Bradley & Corwyn, 2002). Therefore, other relevant measures like home enrichment or parent-child interaction may better identify direct connections with neural maturity. Although we did not find a direct relationship between parental education and network maturation, it would be premature to conclude that this relationship does not exist. For instance, we found a relationship between network maturity and other measures strongly related to parental education, which suggests that parental education impacts cognitive and brain development. Future research should examine multiple dimensions of socioeconomic status to determine if a direct relationship with network maturation can be clarified.

We found that females showed more mature HPC and DMN organization compared to males. Although this result is consistent with stronger within-DMN connectivity in both adult (Ficek-Tani et al., 2023; Ritchie et al., 2018) and youth female populations (Teeuw et al., 2019), there is much less consistency about sex-based differences in the hippocampal network. Studies have reported greater functional connectivity between hippocampal regions in males (Williamson et al., 2024), which may be associated with sex-specific differences in learning strategies that engage the hippocampus (Koss & Frick, 2017; Yagi & Galea, 2019). Structural studies also remain inconclusive, with some reporting no sex differences in hippocampal subregion volumes (Tamnes et al., 2018) and others identifying volumetric differences emerging after certain developmental periods, such as with the onset of puberty (Satterthwaite et al., 2014). Biological factors are linked to brain maturation, likely contributing to our sex-based differences (Gong et al., 2011), however, socialization processes may also play an important role. Girls are often encouraged to engage in sex-biased, socially enriching activities (e.g., pretend doll play), which foster social cognitive abilities like cue processing, perspective taking and theory of mind—functions supported by the DMN (Carter, 2014; Leaper & Lerner, 2015; Lillard, 2017; Hashmi et al., 2020; Kim et al., 2017; Gusnard et al., 2001; Spreng et al., 2010; Witt, 1997). Together, these findings suggest that both biological and experiential factors likely contribute to observed sex differences in HPC and DMN network maturity, though further research is needed to clarify these relationships.

### Limitations and Future Directions

One limitation of this study is sample lack of diversity in parental education level, with a skew toward post-secondary education. This limits our generalizability because low-SES populations are less well represented. A second limitation is the cross-sectional design of the study; longitudinal designs would be valuable to assess dynamic changes in network maturity from childhood to adolescence. A third limitation of the study is the reliance upon parental report measures for adversity. Parent report measurements may suffer from social desirability bias, poor recall, and/or a lack of parental knowledge (Fisher et al., 2011). This is particularly important in the context of quantifying adversity because some experiences may have been overlooked, excluded or understated. Future research should focus on large and diverse samples, including race and racialized experiences, and longitudinal approaches to elucidate the complex influences of environmental factors such as socioeconomic status and adversity on neurocognitive development.

## Conclusion

Overall, the results of our study highlighted the intricate influence that different environmental factors exert on neurocognitive development. Furthermore, it illustrates that while socioeconomic status and early life adversity are connected environmental factors, it is possible to disentangle their influence on development. Our results emphasize that certain environmental characteristics are more impactful on specific areas of development than others. Such findings can help inform targeted interventions that nurture child development by supporting parents’ education and children’s environments.

## Acknowledgements

We would like to thank the Child and Mind Institute for designing and collecting data for the Healthy Brain Network Biobank. We would also like to thank the children, adolescents, and their families for taking the time to participate in studies conducted by the Child and Mind Institute. This work was funded by a Natural Sciences and Engineering Research Council of Canada Discovery grant (RGPIN-2020-05042 to BS RGPIN-2024-06233 to RS and RGPIN-2021-03942 to AS), a CIHR Project Grant (487850 to RS and BS), a Dorothy Killam Fellowship to RS, a SSHRC Insight Grant to RS (435-2017-0936) a SSHRC Insight Grant to RS and BS (435-2024-1375), the University of Western Ontario Faculty Development Research Fund, and a Canadian Foundation for Innovation John R. Evans Leaders Fund to RS (37497). We would also like to thank the Sublime contributions of B<u>. Nowell</u>, E. Wilson, B. Gaugh, and L. Dog.

## Appendix A

### Negative Life Events Scale

|  |
| --- |
| 1. The child's brother/sister was in serious trouble (such as trouble with the law, school, or drugs) |
| 2. The child's close friend had serious troubles, problems, illness, or injury |
| 3. The child suffered from a serious physical illness, injury, or extreme pain (something that required rest of one week in bed, hospitalization, or surgery). |
| 4. The child's brother or sister suffered from serious physical illness, injury, or extreme pain (something that required rest for one week in bed, hospitalization, or surgery). |
| 5. One of the child's brothers or sisters was very angry or upset. |
| 6. People in the child's family (such as his/her parents, brothers or sisters) physically hit each other hard or hurt each other. |
| 7. The child's parent suffered from serious illness, injury, or extreme pain, something that required rest for one week in bed, hospitalization, or surgery. |
| 8. The child's mother or father talked about having serious money troubles (being worried about bills for ordinary things). |
| 9. The child's relatives such as aunts, uncles, grandparents said bad things about his/her mother or father. |
| 10. The child's mother or father fought or argued with his/her relatives such as aunts, uncles, grandparents |
| 11. The child's mother or father acted badly in front of the child's friends (did things like yelled at them or criticized them). |
| 12. The child's mother or father was intoxicated in the child's presence |
| 13. The child's mother or father forgot to do important things for him/her that they promised they would do, such as take him/her on a trip, take him/her to nice places, or come to his/her school or athletic activities. |
| 14. The child's mother or father was arrested or sent to jail. |
| 15. The child's mother or father lost a job |
| 16. A close family member to the child died such as a parent, close uncle, grandparent, or some other relative. |
| 17. A close friend of the child died |
| 18. A close friend of the child moved away |
| 19. The child's father acted very worried, upset, or sad, not because of something the child did. |
| 20. The child's mother acted very worried, upset, or sad, not because of something the child did. |
| 21. The child changed schools. |

