## Supplementary Materials for "Rate of functional network maturity and the role of environmental factors"

**Supplemental Material**

**Executive Function and Social Cognition**

**Parental Education Adversity, Neighbourhood Safety, and Cognitive Ability**

Parental education was positively associated with all WISC measures (see Supp. Figure 1a). As parental education increased, so did participant performance on fluid reasoning (r = 0.24; *p* < 0.001), full-scale intelligence quotient (FSIQ) (r = 0.36; *p* < 0.001), processing speed (r = 0.29; p < 0.001), visual spatial (r = 0.25; *p* < 0.001), verbal comprehension (r = 0.35; *p* < 0.001) and working memory (r = 0.21; *p* < 0.01). Similarly, parental education was negatively associated with social cognition scores (r = -0.24; p < 0.001); children with more educated parents report greater ability (lower scores on SRS scale) to interpret social cues and behaviour (see Supp. Figure 1b).

None of the cognitive variables were significantly associated with adversity (t < 1.64, *p* > 0.10, r < 0.11). Likewise, parental education was not significantly correlated with adversity (r = -0.12; p < 0.07). However, we found neighborhood safety was significantly and negatively correlated with full-scale IQ (r = -0.21; *p* < 0.005), processing speed (r = -0.15; *p* = 0.04), verbal comprehension (r = -0.22; *p* < 0.003), visual spatial ability (r = -0.15; *p* < 0.04), working memory (r = -0.14; *p* < 0.05) and fluid reasoning (r = -0.14; *p* < 0.05) and positively associated with social cognition (r = 0.25; *p* < 0.01), indicating that higher cognitive performance is associated with safer neighbourhood ratings (see Supp. Figure 2a). We also found that the total number of adverse experiences was positively correlated with neighbourhood safety ratings (r = 0.25; *p* < 0.001), suggesting that the more parents reported feeling unsafe in the neighbourhood, the more adversity children experienced (see Supp. Figure 2b). Relatedly, neighbourhood safety was negatively correlated with parental education (r = -0.21; *p* < 0.01), indicating that more educated parents report their neighbourhoods to be safer (higher values for the neighbourhood measure are associated with less safety). All results were FDR corrected for multiple comparison.

**Figure 1**

*Correlation of Parental Education and Cognitive Measures*

*a)*

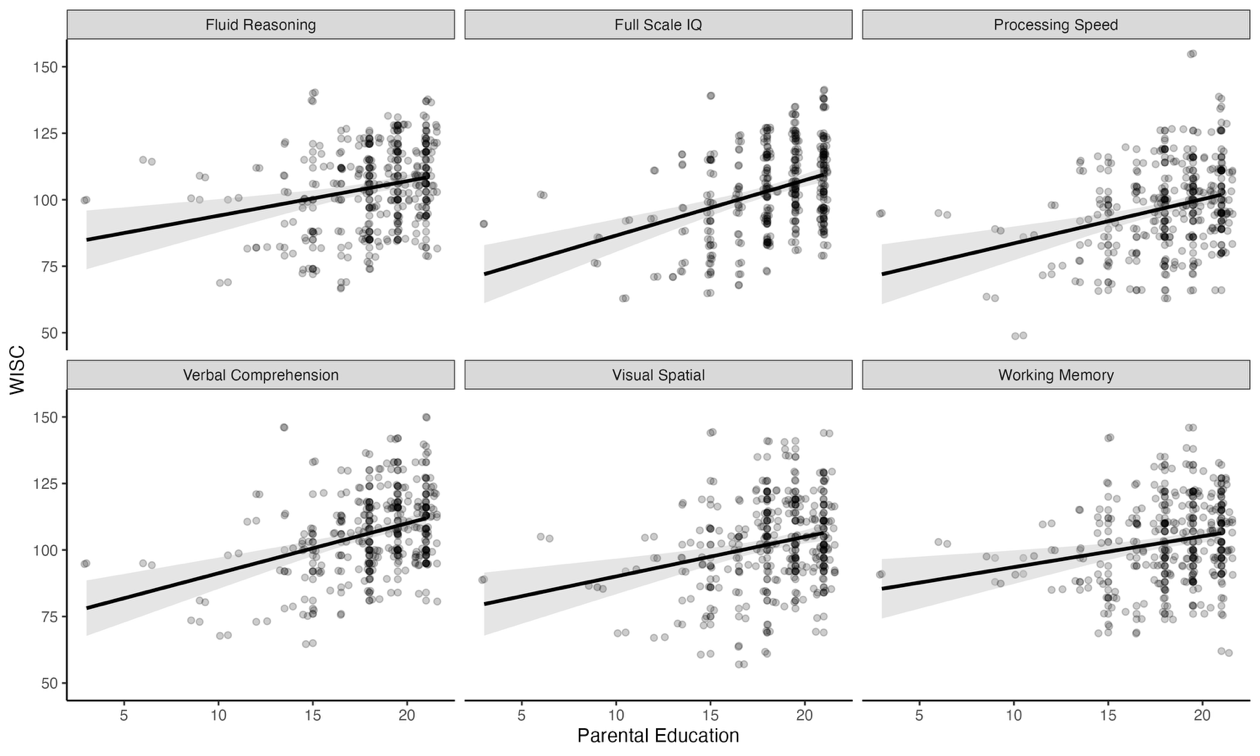

*b)*

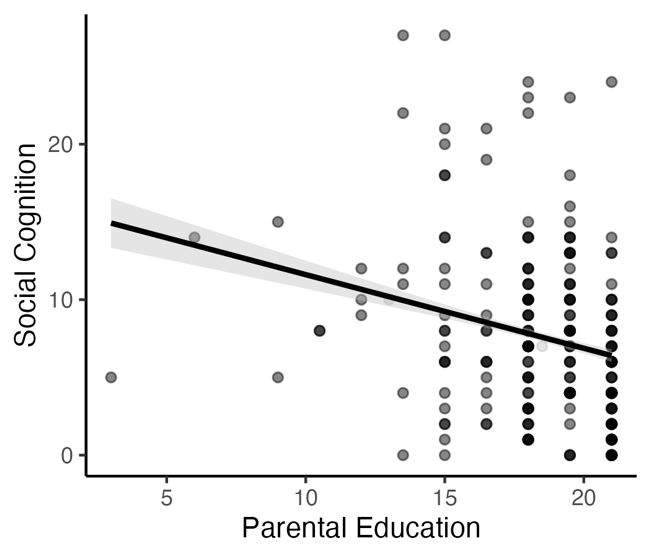

*Note.* Multiple scatter plots representing the relationship between Parental Education and Cognition. Each dot represents a participant's score on the respective cognitive measures, and lines represent the strength and direction of the relationship. a.) Six scatter plots reflecting cognitive performance on fluid reasoning, full-scale intelligence quotient, processing speed, visual spatial, verbal comprehension and working memory, and their respective relationships with parental education. b). A single scatter plot of the relationship between social cognition performance and parental education. Lower scores for social cognition indicate better social cognitive ability.

**Figure 2**

*Correlation of Neighbourhood Safety and Cognitive Measures*

a)

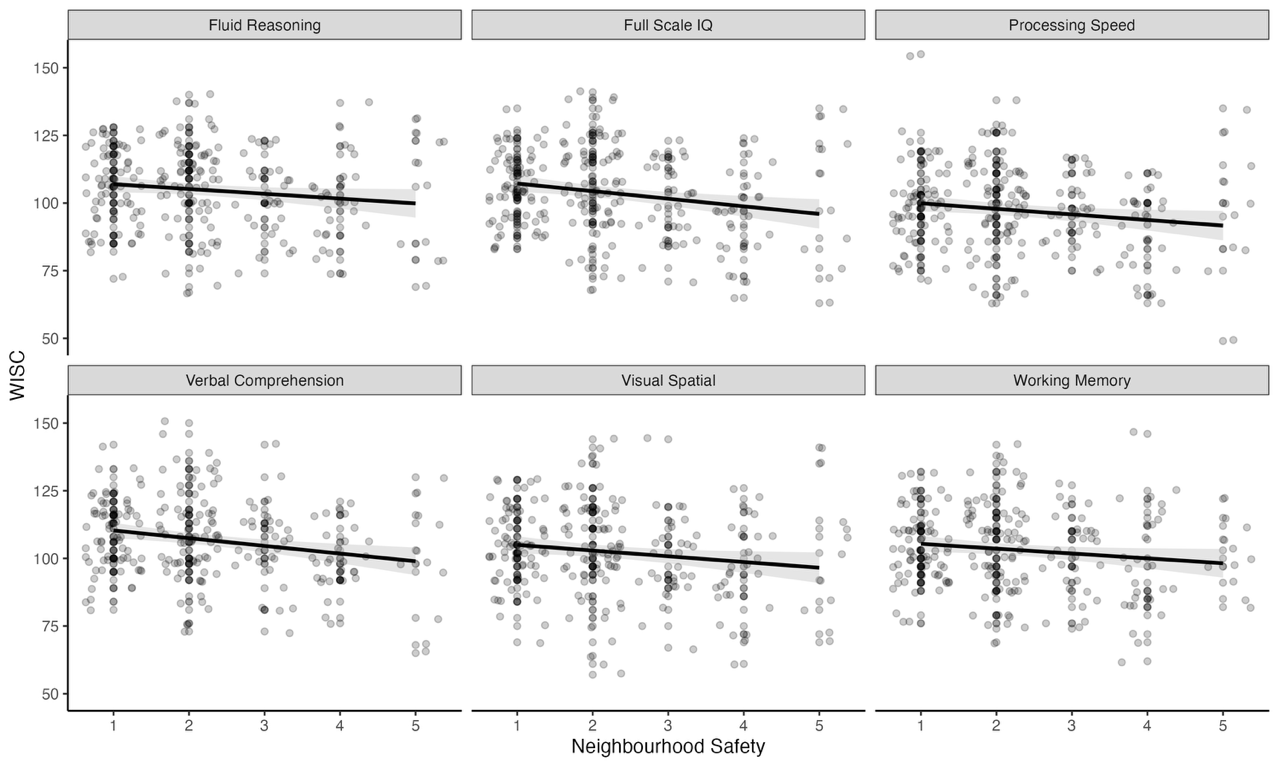

b)

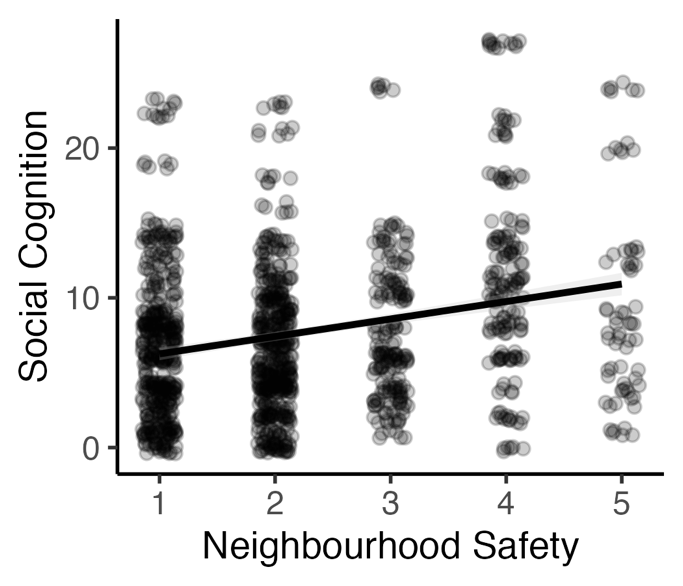

*Note*. Multiple scatter plots representing the relationship between neighbourhood safety and cognition. Each dot represents a participant's score on the respective cognitive measures, and lines represent the strength and direction of the relationship. a.) Six scatter plots reflecting cognitive performance on fluid reasoning, full-scale intelligence quotient, processing speed, visual spatial, verbal comprehension and working memory, and their relationships with ratings of neighbourhood safety. Asterisks represent significant correlations. Higher Neighbourhood Safety values signify neighbourhoods perceived as less safe by parents. b). A single scatter plot of social cognition performance and Neighbourhood Safety. Lower scores for social cognition indicate better social cognitive ability.
